# Predation by *Bdellovibrio bacteriovorus* transforms the landscape and community assembly of bacterial biofilms

**DOI:** 10.1101/2020.08.03.235101

**Authors:** Benjamin R. Wucher, Mennat Elsayed, Daniel E. Kadouri, Carey D. Nadell

## Abstract

The predatory bacterium *Bdellovibrio bacteriovorus* follows a life cycle in which it attaches to the exterior of a Gram-negative prey cell, enters the periplasm, and harvests resources to replicate before lysing the host to find new prey. Predatory bacteria such as this are common in many natural environments, as are groups of matrix-bound clusters of prey cells, termed biofilms. Despite the ubiquity of both predatory bacteria and biofilm-dwelling prey, the interaction between *B. bacteriovorus* and prey cells inside biofilms has received little attention and has not yet been studied at the micrometer scale. Filling this knowledge is critical to understanding the nature of predator-prey interaction in nature. Here we show that *B. bacteriovorus* is able to prey upon biofilms of the pathogen *Vibrio cholerae*, but only up until a critical maturation threshold past which the prey biofilms are protected from their predators. We determine the contribution of matrix secretion and cell-cell packing of the prey biofilm toward this protection mechanism. Our results demonstrate that *B. bacteriovorus* predation in the context of this protection threshold fundamentally transforms the sub-millimeter scale landscape of biofilm growth, as well as the process of community assembly as new potential biofilm residents enter the system.

---

Biofilms are a common mode of microbial life in which cells of one or more species produce surface-attached or free-floating communities that are bound by a self-produced polymer matrix^1–3^. They are thought to be fundamental to microbial ecology in contexts including marine snow carbon cycling^4–6^, the rhizosphere^7^, microbiomes on or within multicellular organisms^8,9^, and acute and chronic infections^10–13^. Biofilm-dwelling bacteria collectively orchestrate their architecture using many mechanisms including the matrix; this architecture then influences surface occupation, dispersal, competition for space and nutrients, and protection from exogenous threats^3,14–16^.

Many studies have shed light on the mechanisms that biofilms use in response to bottom-up selective pressures such as spatial or nutritional competition ^15,17–21^. Others have examined the influence of top-down selective pressures including toxin exposure and predation, which can have a profound impact on the behavior and survival of biofilm communities^16,22–25^. The effects of antibiotic exposure on biofilms have been investigated in detail^26–28^. For example: some but not all antimicrobials are blocked from diffusing completely into biofilms, and those that do permeate biofilms can substantially alter their spatial organization. Other recent work assessed the interaction of bacteriophages and biofilms at single-cell resolution, finding that some biofilms can block phage entry using components of the secreted matrix ^28–30^. The micrometer-scale dynamics of interaction between biofilms and predators that are orders of magnitude larger have received far less attention, however. A key example of such a predator is *Bdellovibrio bacteriovorus*, which is ubiquitous in natural environments^31–35^.

*B. bacterivorous*, a delta-proteobacterium approximately 1 μm in length, most often exhibits an obligate predatory lifestyle in which it targets Gram-negative prey, bores through the outer membrane into the periplasm, harvests cytoplasmic resources to replicate, and lyses the host cell in search of new prey^36–42^. *B. bacteriovorus* has been shown to predate *Escherichia coli* and *Pseudomonas fluorescens* biofilms in static culture and under flow^43^. Numerous studies have isolated *B. bacteriovorus* directly from biofilms on abiotic substrata and the surfaces of animals and plants in aquatic environments^44–49^. Furthermore, predatory bacteria appear capable of navigating spatially complex environments with quite some sophistication; for example, *B. bacteriovorus* can use fungal hyphae to disperse and prey upon distant populations in the soil^50,51^. Predatory bacteria and biofilm communities are thus known to be widespread in nature and commonly to interact ^25,32,39,52,53^, but the details of this interaction have never been studied at single-cell resolution; this is a critical gap in our knowledge of the spatial ecology of *B. bacterivorous* predation.

In aquatic environments, predatory bacteria are strong modulators of the *Vibrio* clade^53^, and *V. cholerae* is a known susceptible prey target to *B. bacterivorous* in estuarine environments^33,52^. We therefore chose *Vibrio cholerae* as a model organism to examine *B. bacteriovorus* interaction with prey biofilms, because its architectural dynamics and matrix components have been characterized in depth^54–56^. Using a combination of microfluidic culture, confocal imaging, and detailed spatial analysis, we explore how biofilm structure and composition can affect the outcome of bacterial predation pressure, as well as the broader ecological impacts that predation can have on a biofilm community. We find that exposure to bacterial predators fundamentally alters the landscape of biofilm growth and communal defense against infiltration by newly arriving planktonic bacteria.

## Results

### *V. cholerae* biofilms have a maturation threshold for protection from *B. bacterivorous*

To evaluate the interaction between pre-formed resident *V. cholerae* biofilms and their bacterial predators, we first cultivated *V. cholerae* on glass surfaces in microfluidic devices (see Materials and Methods). Approximately 48h after the initial surface inoculation and initiation of flow, we introduced *B. bacteriovorus* into the chambers over a period of 30 min (2.5×10^9^ PFU/mL at 0.2μL/min flow rate, or approximately 1.5×10^7^ *B. bacteriovorus* cells in total), followed by resumption of predator-free medium flow for the remainder of the experiment. Biofilms were then imaged through their entire 3D volume by confocal microscopy.

Successful predation and bdelloplast formation could be seen throughout the microfluidic arena among singleton prey *V. cholerae*. Cells on the periphery of biofilm clusters appeared susceptible as well, but the centers of larger biofilm clusters remained devoid of predator cells (Figure 1A). It is possible that protection of cells in the interior might be temporary, and that over time *B. bacteriovorus* could mobilize and consume cells throughout the biofilm. However, this was not the case: images taken 48 h after initial predator exposure showed that cells on the interior of these clusters remained unexposed to predation; remaining *B. bacteriovorus* cells were immobilized in the matrix milieu around resident prey throughout the expanding front (Figure 1B). These results suggest that one or more features of *V. cholerae* biofilm architecture might inhibit predator cells from penetrating the interior of the biofilm after initial attachment.

**Figure 1.**
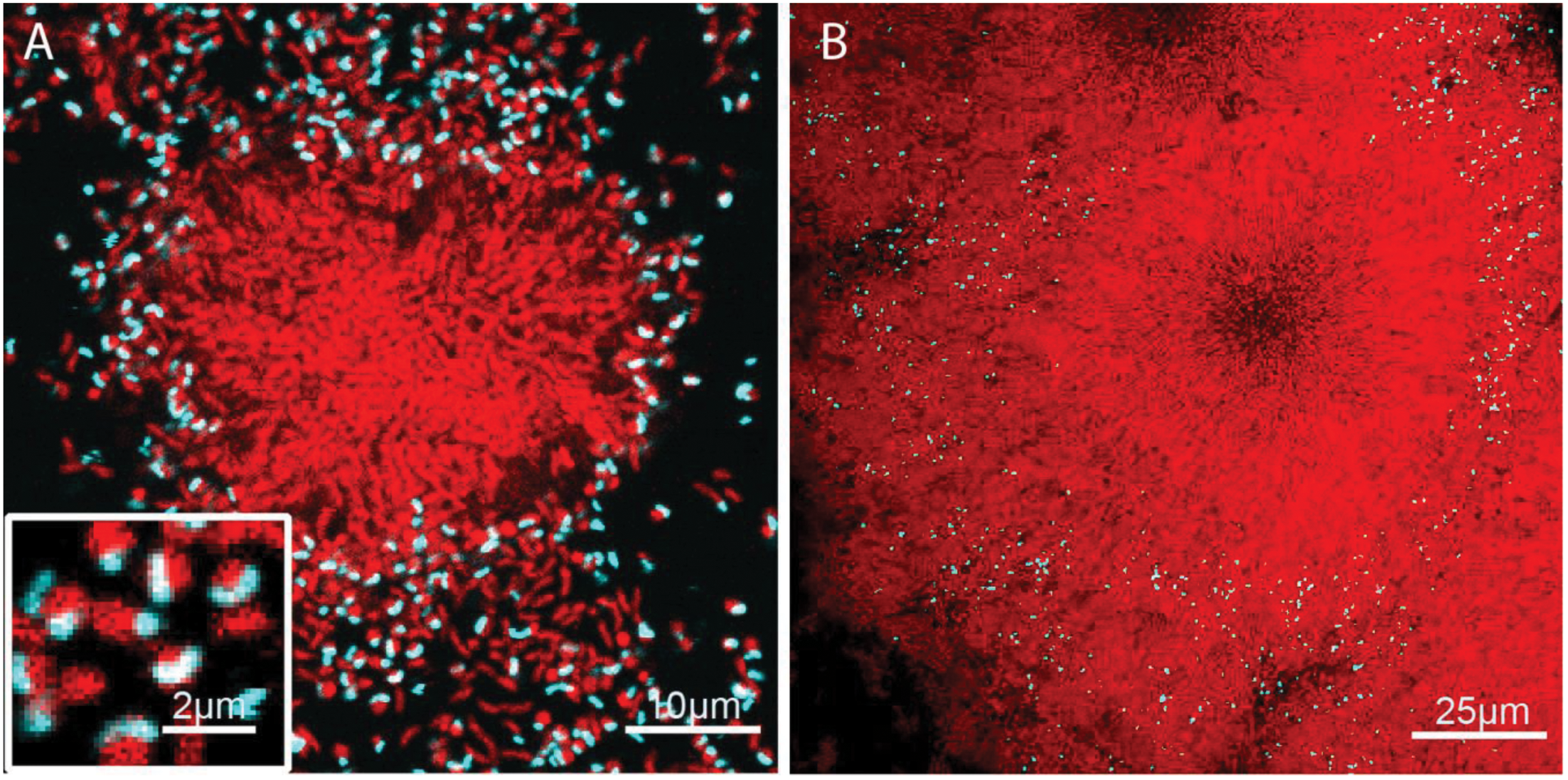
An illustration *V. cholerae* biofilm clusters following *B. bacteriovorus* exposure. Prey biofilms (red) were grown for 48 h prior to exposure to predator cells (cyan). (A) 30 min after introduction, predator cells have preyed upon singleton cells, forming bdelloplasts (inset). Predator cells also appear able to access hosts on the periphery, but not within the innermost regions, of *V. cholerae* host biofilm clusters. (B) 48 hours after introduction, *V. cholerae* demonstrates net positive growth, trapping *B. bacteriovorus* in the expanding front.

We next sought to understand what components of *V. cholerae* biofilm structure influence spatial access by predatory cells. Prior work has linked protection of biofilms from entry by bacteriophages and competing microbes to the production of proteinaceous or polysaccharide constituents of the biofilm matrix^16,21,28^. Following this precedent, we were curious as to the contribution of the matrix in protection from *B. bacteriovorus* predation. To pursue this question we introduced a 3x-FLAG epitope to the N-terminus of the *V. cholerae* matrix protein RbmA; this construct allowed us to directly visualize the matrix without altering its function^21^. RbmA has been extensively characterized as a key matrix component, along with vibrio polysaccharide (VPS), in controlling cell-cell packing and alignment architecture within biofilms of this species^3,14,54,57^. Our visualizations showed that *B. bacterivorous* localized within the outermost layers of cells and matrix material in the periphery of larger biofilm clusters. *V. cholerae* cells outside of the matrix were frequently preyed upon (Figure 2A-B). Visual inspection alone could not determine whether or not proximity to matrix was sufficient on its own to prevent prey killing by predatory bacteria, as is the case in protection of *E. coli* against attack by T7 bacteriophages^16,29^.

**Figure 2.**
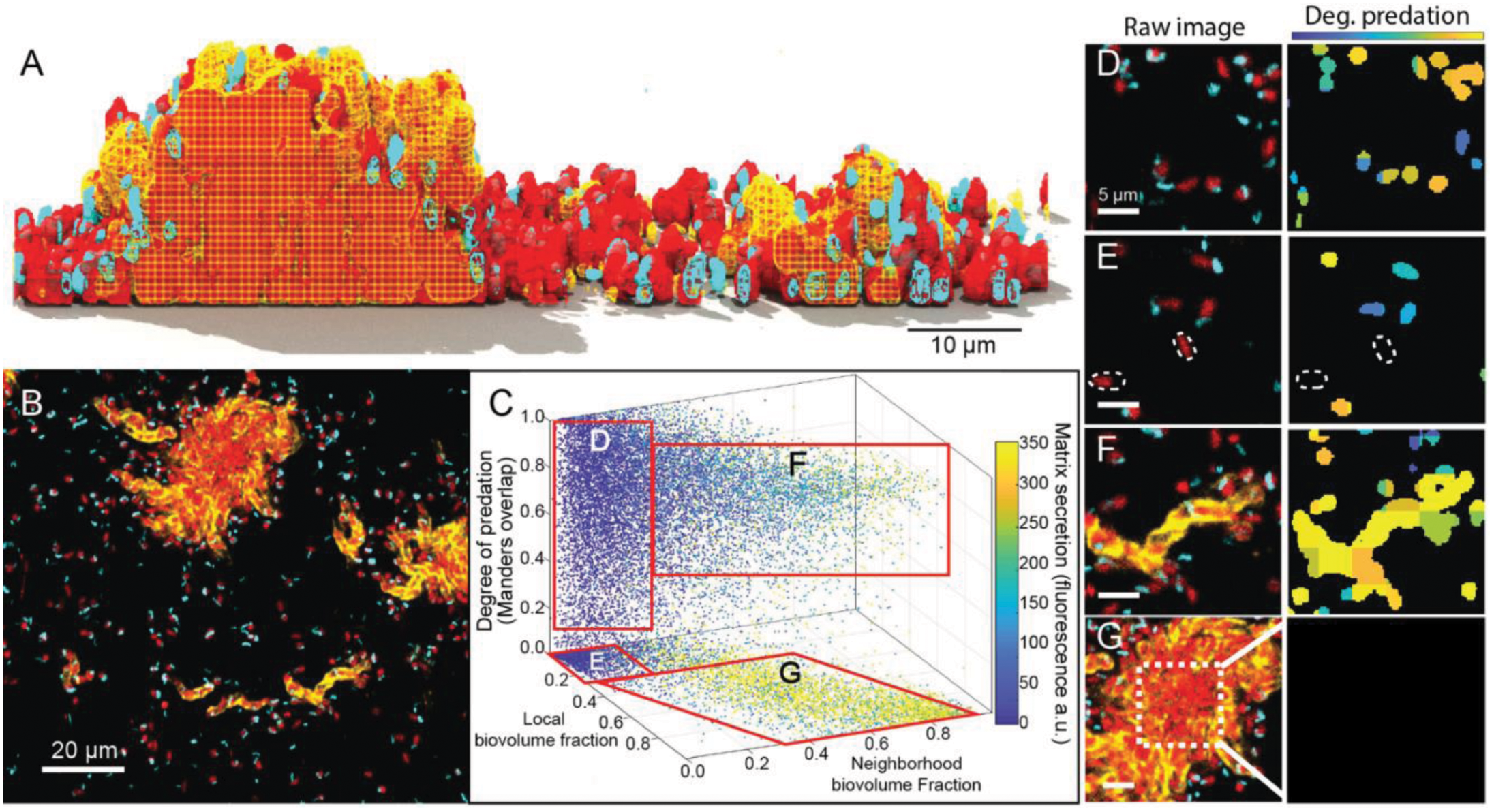
Image cytometry analysis of *V. cholerae* biofilm matrix secretion, cell packing architecture, and susceptibility to *B. bacteriovorus* predation. (A) A 3-D rendering of *V. cholerae* biofilms (red), secreted matrix (yellow), and *B. bacteriovorus* (cyan) showing a vertical cross section of the biofilm 2 hours after the introduction of predators. The matrix is rendered as a mesh to help visualize the embedded cells. (B) Raw fluorescence image showing a horizontal cross section of the matrix-labeled biofilm (same color scheme and timepoint as in panel A). (C) Image analysis of biofilms exposed to predatory bacteria after 2 hours. The X and Y axes denote local and neighborhood biovolume fraction, respectively. The vertical axis denotes the degree of predation. Any points off the bottom plane denote host cells in the process of being killed by predatory bacteria. Data points are color-coded according to local matrix fluorescence intensity. (D-G) Raw images and corresponding heat maps for degree of predation. In the raw images at left, host cells are red, predators are cyan, and matrix is yellow. In the heatmaps at right, blue/teal indicates a predator cell attached to a host cell, and orange/yellow indicates a predator cell is inside the host. (D) isolated singleton cells are fully exposed and tend to be killed off by *B. bacteriovorus*, though some singleton cells have not yet been found by a predator, highlighted by the dotted outlines in (E). (F) Small biofilm clusters that are producing extracellular matrix are nevertheless fully susceptible to predation. (G) Though the periphery regions of large biofilm clusters are still susceptible to predation – as in (F) – the internal regions of these clusters with high cell-packing are fully protected.

To resolve this uncertainty, sought to measure at spatial high resolution the amount of secreted matrix, the cell-cell packing density among prey *V. cholerae* cells, and the relationship between these biofilm architecture features and the extent of local predation by *B. bacteriovorus*. To accomplish this, we used the BiofilmQ analysis framework to segment predator and prey biovolumes and to dissect them into a 3-D grid, with each cubic grid unit measuring 2.6 µm on a side. At this resolution, the grid units could contain 3-5 cells of *V. cholerae* and/or *B bacteriovorus*. For each grid unit we calculated i) the local matrix accumulation around *V. cholerae*; ii) the local biovolume fraction (i.e. how much of each grid unit was occupied by *V. cholerae*); iii) the neighborhood biovolume fraction (i.e., how much of a 10 µm diameter bubble around each unit was occupied by *V. cholerae*); and finally iv) an overlap coefficient between *V. cholerae* and *B. bacteriovorus* (which corresponds to the degree of predation, see Materials and Methods). Note that the local and neighborhood biovolume fractions are both proxies for cell-cell packing of prey *V. cholerae*, but on two spatial scales, and so they yield different information about localized versus surrounding cell-packing architecture. For example, a small biofilm cluster of 5-10 cells that have begun to produce matrix typically has high local volume fraction, because its cells are all in close proximity; but such a nascent biofilm also has low neighborhood volume fraction, because it has not yet expanded into a mature biofilm cluster. Visual representations of the segmentation process and the parameters we calculated can be found in SI Figure S1.

Using the metrics described above we analyzed *n* = 23 independent image stacks (summarized in Figure 2C), which revealed four different biofilm sub-populations. We label these D-G for correspondence with examples of each in panels D-G of Figure 1. Population D includes singleton *V. cholerae* cells with zero matrix, low local and neighborhood biovolume fractions, and which have been preyed upon by *B. bacteriovorus* (Figure 2D). Population E includes singletons much like population E, but which have not yet been found by a predator cell (Figure 2E). Population F includes *V. cholerae* clusters that have begun producing matrix, but which had not yet formed hemi-spherical groups; this sub-population had detectable matrix signal, high local biovolume fraction, but low neighborhood biovolume fraction (Figure 2F). Also in group F were units on the outer periphery of larger biofilm clusters. These cells, despite accumulating matrix and high local density, were susceptible to predation (SI Figure S2). Lastly, population G included groups of cells on the interior of larger biofilm clusters; these had high matrix accumulation, high local and neighborhood biovolume fractions, and almost complete protection from predation (Figure 2G). Overall, these results suggest that local matrix accumulation alone is not sufficient for protection from *B. bacteriovorus*; rather, a combination of matrix secretion and cell-cell packing is at play.

To further explore the interaction between matrix production, cell-cell packing, and predation protection, we studied two additional mutants and their susceptibility to *B. bacteriovorus*. One is a *vpv*^W240R^ point mutant that constitutively produces extracellular matrix – we refer to this strain as a matrix hyper-secretor. The other, Δ*rbmA*, harbors a clean deletion of the *rbmA* locus and therefore cannot produce the core matrix protein RbmA. The hyper-secretor rapidly generates highly compact biofilm clusters relative to wild type^58–60^, and Δ*rbmA* produces biofilms with far looser cell-cell packing and altered cell orientation architecture^3,14,19,21,55^. These strains – and WT for comparison – were grown in monoculture microfluidic devices and subjected to a single dose of *B. bacterivorous* (Figure 3A-C).

**Figure 3.**
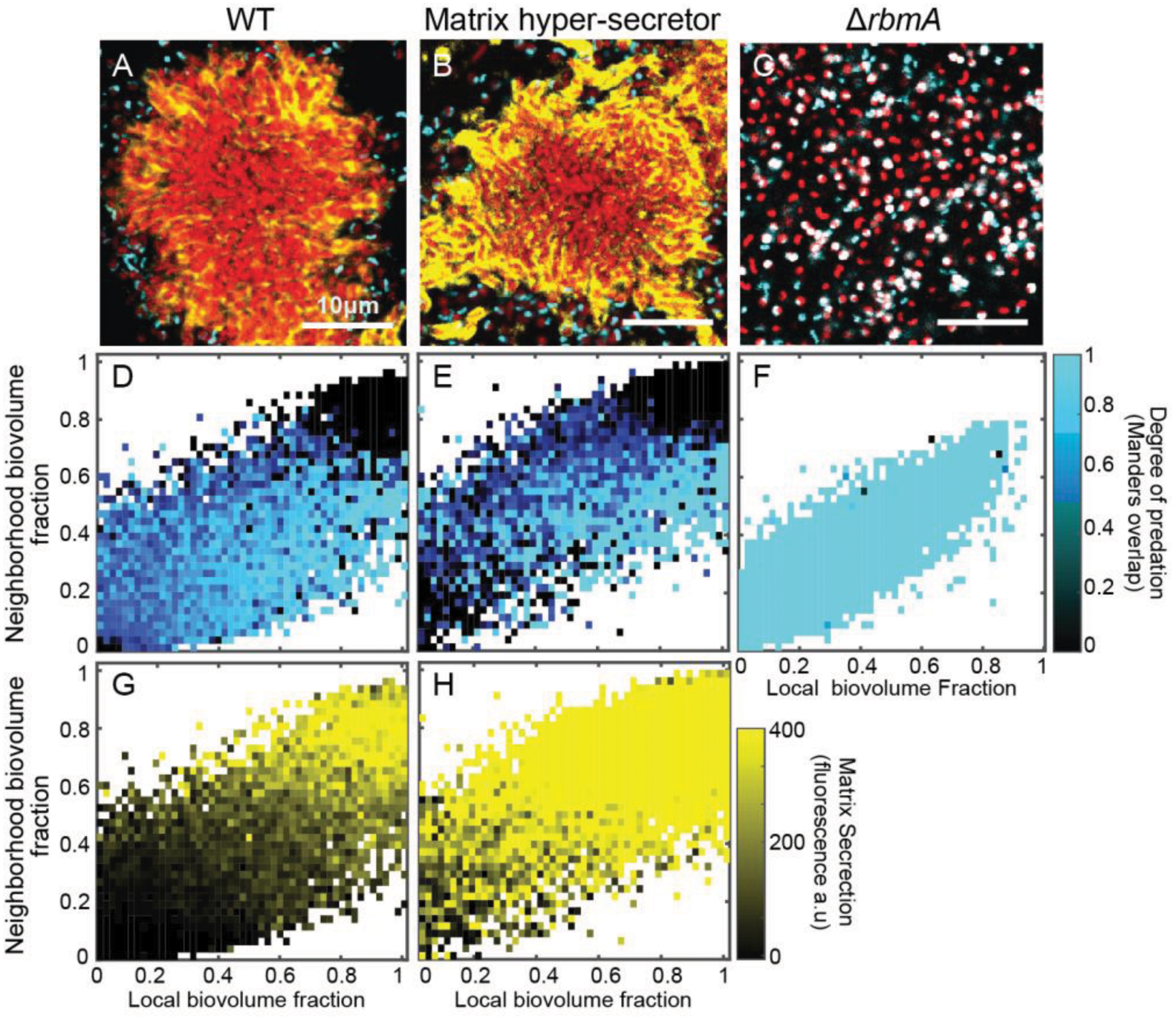
A critical threshold of neighbohood biovolume fraction correlates with host cell protection from predation. (A-C) Images of *V. cholerae* biofilm clusters of wild type, matrix hyper-secreting, and Δ*rbmA* strains 2 hours after predator introduction. *V. cholerae* cells are shown in red, immunostained RbmA-FLAG is shown in yellow, and *B. bacteriovorus* is shown in cyan. Biofilms were segmented and analyzed by dissecting the total system into a cubic grid as detailed in the main text. The segmented biovolumes in each grid are analyzed individually to produce the kymographs described below. (D-F) Heatmap plots for the degree of predation in biofilms of the three strains shown in (A-C), respectively. The horizontal axis denotes local biovolume fraction, and the vertical axis denotes neighborhood biovolume fraction. Black squares correspond to biofilm volume units that are protected from predation; dark blue squares denote areas with predation initiating at the cell exterior; and light blue squares denote areas fully predated. Note the critical threshold neighbohood volume fraction of approximate 0.8 past which biofilms are protected from predation. (G-H) Heatmaps plots for the degree of matrix accumulation in biofilms of the two strains shown in (A-B), respectively. There is no entry for the Δ*rbmA* strain, because it cannot produce the matrix protein being immunostained. Axes are as for (D-F). The black-to-yellow scaling relates the matrix accumulation for each point. Note in comparing (E) and (H) in particular that high matrix production by itself does not confer predator protection; rather matrix-replete regions of the biofilm must first reach the critical neighborhood cell packing threshold before predators can be exluded.

The resulting image data were again segmented and dissected into a cubic grid for spatial analysis as described above. Panels D-F in Figure 3 show heatmaps of local versus neighborhood biovolume fraction with points color-coded according to predation state; panels G-H in Figure 3 show analogous heatmaps, but with points color-coded according to local RbmA accumulation. From this analysis it is evident that both WT and matrix hyper-secreting strains have a critical neighborhood biovolume fraction (∼ 0.8) past which patches of cells are protected from predator exposure (Figure 3D-E; SI Figure S3). Cell clusters of the matrix hyper-secreting strain reached this threshold more quickly, and so had greater total protection against predation (SI Figure S4, S5). Importantly, however, even though the matrix hyper-secreting strain has a higher signature of matrix secretion (Figure 3G-H), its threshold biovolume fraction for protection against *B. bacteriovorus* was the same as that of WT. By comparison, biofilms of the Δ*rbmA* strain never reach the biovolume fraction threshold required for protection against predator attack, and nearly all cells are killed (Figure 3F).

Altogether these data suggest that is not the extracellular matrix on its own but rather the collective cell-cell packing that emerges from cell-matrix and cell-cell interaction that ultimately provide protection against predation by *B. bacteriovorus*. Another striking implication of our analysis is that there is not one but two advancing fronts on the outer periphery of growing *V. cholerae* biofilms. The first is the true outer layer of biofilm expansion in which cells are producing extracellular matrix but have not yet achieved the cell-packing required for *B. bacteriovorus* protection. The second front, lagging behind the first, is that at which matrix and cell-packing have consolidated, conferring lasting protection against invasion by bacterial predators. Our results imply that the rate of consolidation of this secondary front exceeds the rate of infiltration and predation by *B. bacteriovorus* on the biofilm periphery, allowing the biofilm to maintain positive net growth despite grazing by the predator population in the outermost biofilm layer.

### *B. bacterivorous* predation transforms the landscape of *V. cholerae* biofilm growth

Our results thus far establish a critical cell-packing threshold past which biofilms of *V. cholerae* survive exposure to *B. bacteriovorus* (Figure 3D-E; SI Figure S3); though the predator can continue grazing on the outer periphery of these biofilms, the prey cell clusters maintain positive net growth. There are precedents for this observation, but at much larger spatial scales in the context of forest ecology. Our findings are analogous to browsing and fire traps well known to limit the recruitment of tree saplings to adult trees – only saplings past a size threshold survive herbivore grazing and fire to become adult trees^61,62^. Depending on grazing and fire frequency, this effect can generate vastly different distributions of tree biomass distribution on continental scales^63^. With this analogy in mind we were curious as to the impact of predation on biofilm distribution: how does exposure to *B. bacteriovorus* influence the sub-millimeter scale landscape of *V. cholerae* biofilms?

We explored this question by repeating the experiment above with a different imaging regime. *V. cholerae* was grown microfluidic devices for 48 h before a single introduction of *B. bacterivorous*, followed by a return to predator-free media influx. In control treatments, the same tubing exchanges were performed, but no predators were introduced. We then imaged the biofilms by confocal microscopy 48 h later, which revealed dramatic differences between the two treatments. Control chambers contained a wide distribution of cell cluster sizes (Figure 4A). The frequency distribution of neighborhood biovolume fraction in this condition was broad with a shallow peak at 0.5 (Figure 4C).

**Figure 4.**
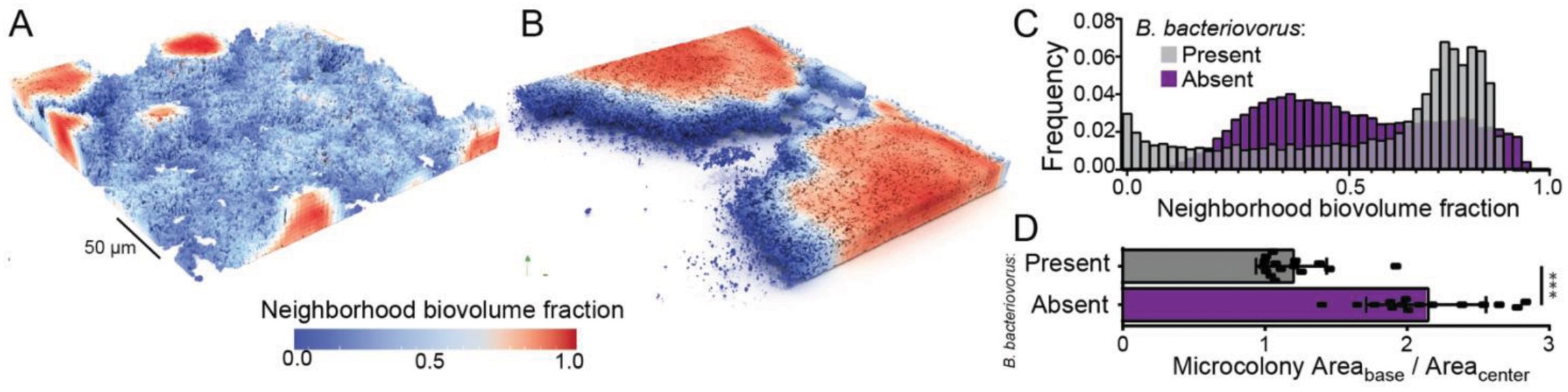
Exposure to predation by *B. bacteriovorus* shifts the microscopic landscape of host biofilms. A) In the absence of predatory bacteria, *V. cholerae* produces biofilms with abundant small clusters with high internal neighbor volume fraction and low peripheral neighborhood volume fraction. B) Under predation by *B. bacteriovorus*, single cells and small colonies below a neighborhood cell-packing threshold are exposed and killed, leaving few remaining clusters which are then free to grow very large. C) Frequency distributions of neighborhood volume fraction for biofilms exposed or unexposed to *B. bacteriovorus* predation. Biofilms with predators present show a strong shift toward high neighborhood volume fraction. D) Quantification of the average ratio of basal area to mid-plane area for biofilms with and without exposure to predators. Exposed biofilms, because they have room to grow into much larger columnar structures, have a ratio of ∼1; while in unexposed biofilms, clusters compete more for space and remain semispherical, such they are larger at their base than they are at their mid-plane.

Biofilms exposed to *B. bacteriovorus* were strongly shifted toward very large cell clusters that had reached the ceiling of the chambers and grown into columnar structures, in contrast to the hemispherical biofilm clusters observed in the control chambers (Figure 4B). We could test whether the difference in biofilm cluster shape between the two treatments was consistent across all replicates by measuring the ratio of biomass at the base of biofilm clusters to that at the chamber mid-plane. This ratio was ∼2 in control chambers but transitioned to 1 in predator-exposed chambers, reflecting the change from hemispherical to columnar cell groups (Figure 4D). The distribution of neighborhood volume fraction for predator-exposed chambers showed a pronounced shift toward high values in the range of 0.8, the critical cutoff identified above for protection from predator attack (Figure 4C). This shift occurred quickly, within the first 16 hours after predator exposure (SI Figure S6). In chambers with predators introduced, the space around large clusters was mostly unoccupied, presumably due to killing by *B. bacteriovorus*, which contrasted sharply with control chambers in which areas surrounding cell clusters were occupied by nascent biofilm clusters or cell monolayers (SI Figure S7).

### *B. bacterivorous* exposure alters biofilm surface structure and allows infiltration by newly arriving bacteria

An additional observation from our long-term imaging experiments was that among biofilm clusters which survive predator-exposure, their outermost layers – which remained susceptible to *B. bacteriovorus* – look to be more loosely packed and porous than those of biofilms in the control condition (Figure 4A-B). Cell packing in the exterior of biofilms is an important element of a community barrier function in *V. cholerae* and other microbes, which protects against intra- and inter-specific infiltration ^21,28^. Typically, *V. cholerae* biofilms rarely allow for successful surface colonization by other bacteria, and they are extremely resistant to enter into their interior^21,28^. The packing architecture that confers this protection is a result of cell-matrix and cell-cell interactions which altogether form the basis of structural strength in their biofilms. We hypothesized that by killing a fraction of cells on the biofilm exterior layer, *B. bacteriovorus* partially compromises this packing architecture, perhaps rendering them less resistant to entry by other bacteria including conspecific competitors. To test this idea, we once again grew *V. cholerae* biofilms for 48 hours and subjected them to a single dose of *B. bacteriovorus*. 48 hours later, we introduced new competitors to the environment in the form of an isogenic *V. cholerae* strain that produced a different fluorescent protein than the resident biofilm, so the two could be distinguished from each other and the predatory cells.

In control chambers without predator exposure, resident biofilms blocked invasion of newly introduced cells: as seen previously^21^, few invaders could be found on the biofilm outer surface, and none made it into the biofilm interior (Figure 5A, D). In contrast, predator-exposed biofilms permitted substantial infiltration of competitors past their outer boundaries (Figure 5B-D). Quantifying these results by image analysis, invasion of competitors into predator-exposed biofilms was 40-fold greater than that of control biofilms (Figure 5E). Areas of resident biofilms with many *B. bacterivorous* cells present also appeared to have a high density of invading cells (Figure 5C,D). To quantify this observation, we measured the localized signal intensity of invading cells surrounding resident cells and compared this metric with the localized degree of predation by *B. bacteriovorus*. We found a linear correlation between the number of invading cells present in a given area as a function of how much predation that area had experienced (Figure 5F). This outcome is consistent with our hypothesis that *B. bacteriovorus* predation disrupts local biofilm architecture and renders it more openly exposed to entry by other cells. In this respect *B. bacteriovorus* not only alters the structure of the outermost biofilm front but also fundamentally changes the ecology of biofilm assembly as new and potentially competing (but-non-predatory) cells enter the system.

**Figure 5.**
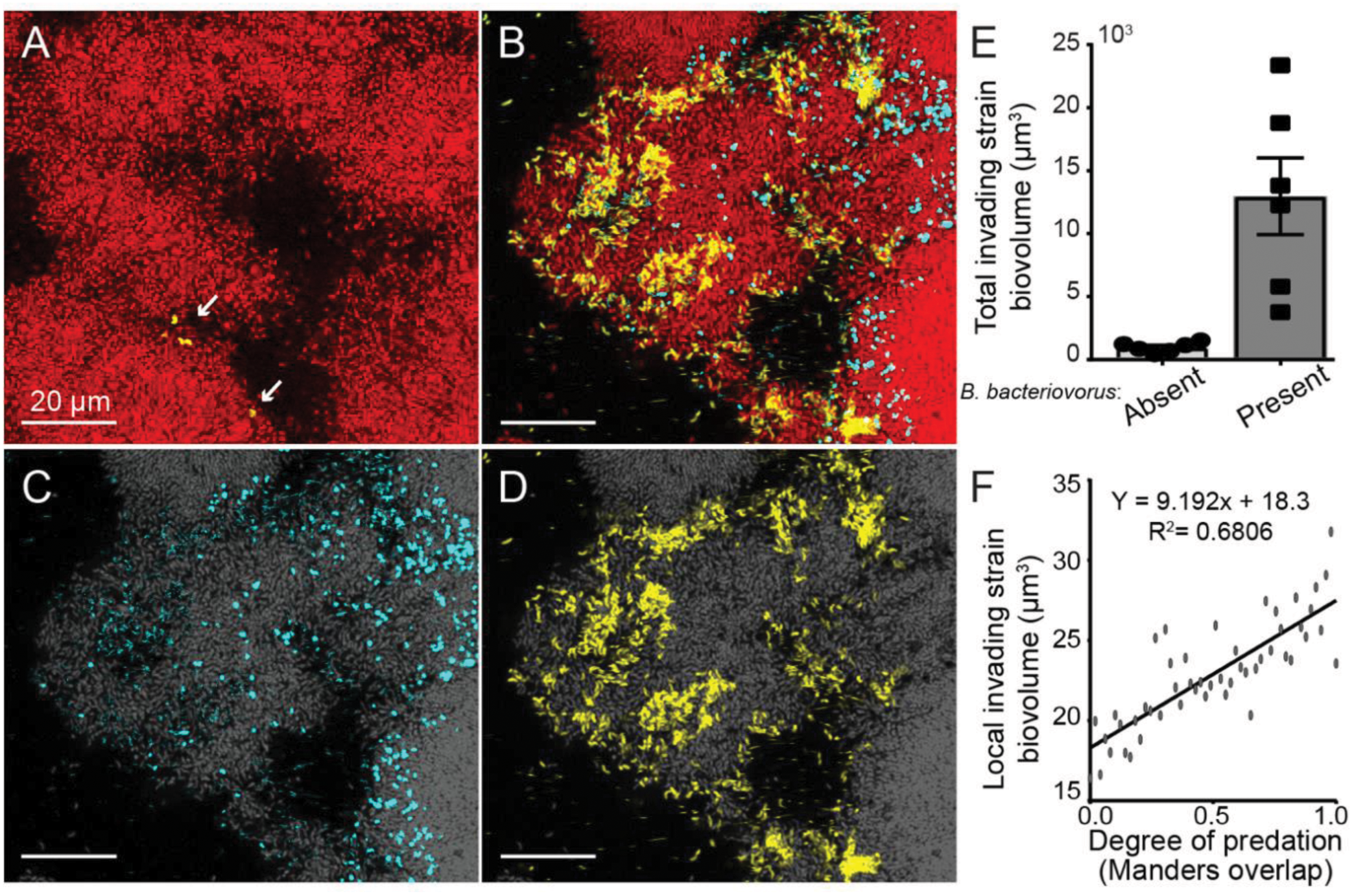
*B. bacteriovorus* exposure on the periphery of *V. cholerae* biofilm clusters renders them susceptible to infiltration by other bacteria. (A) In the absence of predator exposure, *V. cholerae* biofilms are highly resistant to invasion by conspecific cells. The resident biofilm is shown in red, and invading cells are shown in yellow. (B) Resident biofilms that have been exposed to predation by *B. bacteriovorus* (blue) have a more loosely structured periphery, and as a result, invading conspecifics are able to enter well past the outer boundary of the resident biofilm. (C) Image of the predator bacteria (blue) and (D) invading conspecific cells (yellow) distributed in the outer resident biofilm layers (resident biofilm in grey). (E) Measurement of the differences in total invading cell biovolume across the whole biofilm, in the absence or presence of *B. bacteriovorus*. (F) Within biofilms exposed to predation, the degree of invasion within any given local area scales linearly with the degree of predation in that area.

## Discussion

Predator-prey interactions in the context of microbial biofilms are almost certainly widespread in nature; we are only in the early stages of understanding the micrometer-scale processes that determine the outcome of these encounters, the underlying molecular mechanisms of these encounters, and the consequences for microbial ecology and evolution. Major steps forward have recently been made to understand phage-biofilm interaction^16,28,64,65^, and landmark papers have begun to characterize predation by larger protist predators and cells of metazoan immune systems ^23,66–68^ at high resolution. Biofilm grazing by metazoans has been studied, but primarily at macroscopic scales^69–71^. *B. bacteriovorus*, a ubiquitous threat to prey bacteria, has been investigated interacting with biofilms, but again primarily via macroscopic assays^43,52^. Here we build on this foundation with the first single-cell resolution live imaging of *B. bacteriovorus* preying upon biofilms of *V. cholerae*. The *V. cholerae* cell-cell packing threshold that we discovered, past which predators are not able to access their prey, reveals novel insights into the mechanisms of architecture maturation, and it leads to fundamental transformations of biofilm structure and community assembly.

We hope to have demonstrated that high-resolution imaging and analysis of predator-prey interactions inside biofilms is a critical area open for further investigation in light of the enormous diversity of predator types and biofilm structure, and their potential influence on each other. Prior work has intimated, for example, that *B. bacteriovorus* is able to kill whole biofilms of *E. coli* and *P. aeruginosa*, even after the prey have produced relatively large cell groups^43,52^. Understanding the key determinants of successful predation of prey biofilms will be important for our knowledge of the natural history of biofilm formation in different species, and also for the potential uses of *B. bacteriovorus* as an antimicrobial therapeutic.

## Author Contributions

CDN and DEK conceived the project. CDN supervised the project. CDN and BRW designed experiments. BRW performed experiments and image processing of microscopy data. CDN and BRW analyzed data. DEK and MS provided reagents/tools. CDN and BRW wrote the paper with input from DEK.

## Acknowledgements

We thank R.W. Baker, A. Persat, N. Rigel, B. Ross, D. Schultz, and members of the Nadell Lab for their comments and suggestions on earlier versions of this manuscript. BRW is supported by a Gilman Fellowship from the Department of Biological Sciences at Dartmouth. CDN is supported by NSF grant MCB 1817342, NSF grant IOS 2017879, a Burke Award from Dartmouth, NIH grant 2R01AI081838 to PI Robert Cramer, NIH grant P20-GM113132 to the Dartmouth BioMT COBRE, and grant RGY0077/2020 from the Human Frontier Science Foundation with co-PI A. Persat.

## Supplementary Information

### Supplemental Materials and Methods

#### Strains and media

The full strain and plasmid list for this study can be found in **Table S1**. Prior to experiments, *V. cholerae* strains were grown overnight in lysogeny broth (LB) in a shaking incubator. *B. bacteriovorus* stocks were obtained via co-culture with prey and subsequent filtering. A full outline of the methods has been described previously^1^. Modifications to *V. cholerae* were made using *E. coli* strain S-17-λpir carrying the allelic exchange vector pBW1 as previously described^2^. Antibiotics and reagents used for counter selection were the following concentrations: 100µg/ml ampicillin, 50µg/ml kanamycin, 50µg/ml polymyxin B, 5% sucrose. All reagents were obtained from Millipore sigma unless otherwise stated.

#### Microfluidic assembly

Poly-dimethysiloxane (PDMS) was used to cast microfluidic chambers using standard soft lithography techniques^3,4^. The chambers were bonded to #1.5 coverslips measuring 36mm by 60 mm (WxL). The chambers used for this study had dimensions of 3000µm x 500µm x 75µm (LxWxD). In order to run media through these chambers 1ml of M9 with 0.5% glucose was loaded into 1mL BD plastic syringes. 25 gauge needles were affixed to the syringes and #30 Cole Palmer PTFE tubing with an inner diameter of 0.3mm was placed over the end of the needle. The other end of this tubing was then placed into pre-bored holes in the microfluidic device. An additional length of tubing was run from the auxiliary channels in the device to a vacuum line, thereby preventing bubbles from entering the system. Syringes were mounted Pico Plus Syringe Pumps (Harvard Apparatus)

#### Biofilm growth conditions and matrix staining

Biofilms were grown in the microfluidic chambers described above. Overnight cultures of *V. cholerae* were back-diluted into M9 minimal medium with 0.5% glucose and allowed to re-enter exponential phase (OD_600_ = 1.0) to acclimate to the media conditions used for biofilm growth (i.e. M9 minimal media with 0.5% glucose). These cultures were inoculated into chambers without flow to allow surface colonization for 1 h. After this period a flow rate of 0.2µL/min was established for the remainder of the experiment. All experiments were performed at room temperature. For matrix straining experiments in which *V. cholerae* harbored C-terminal fusion of 3xFLAG to RbmA, an anti-FLAG antibody conjugated to a Cy3 fluorophore added to the influx medium (M9 minimal with 0.5% glucose) at 1µg/ml.

#### Introduction of predators and invaders

Introduction of predators was done in a similar fashion to the chamber colonization of *V. cholerae. B. bacteriovorus* cultures were diluted to an OD600 of 0.5 (2.5 x10^9^ PFU/mL) before being inoculated into the chamber. To do this, the media tubing was briefly removed, and the *B. bacteriovorus* was inoculated into the culture via a micropipette. The media tubing was then returned to its position, and flow was resumed 30 minutes after introduction of predators. For experiments in which biofilms were challenged with conspecific invading *V. cholerae*, a similar regime was carried out. Overnight culture of *V. cholerae* housing a different fluorescent protein than the resident biofilms was diluted to an OD600 of 1 and then inoculated into the chambers. Tubing was replaced and flow was resumed 30 minutes after inoculation.

#### Microscopy

Imaging of the biofilms was performed with a Zeiss LSM 880 laser scanning confocal microscope. The microscope used either a 40x /1.2 N.A. water objective or a 10x/ 0.4 N.A. water objective. A 488-nm laser line was used to excite the GFP contained in the *B. bacteriovorus*. To Image *V. cholorae*, a 594-nm laser was used to excite mKate2 in the resident strain and a 543-nm laser was used to excite mKO-κ for the invading strain. This 543-nm laser was also used to excite the Cy-3 Fluorophore on the anti-FLAG antibody for matrix staining.

#### Image Analysis

To obtain data for image analysis, several image stacks were taken at independent locations within each chamber. These image stacks were then analyzed using the framework BiofilmQ. A detailed explanation of BiofilmQ can be found in several previous studies^5,6^.

#### 3D renderings

3D renderings were created by first using the VTK output feature present in BiofilmQ. These files could then be processed in ParaView and rendered using Osprey ray tracing.

#### Statistics

Statistical analyses were performed in GraphPad prism. All pairwise comparisons were made using Wilcoxson signed ranks test with Bonferroni correction. Differences between frequency distributions were compared via Kolmogorov-Simirnov tests.

**Table S1.** Strains and plasmids.

| Strain | Relevant markers/ Genotypes | Source |
| --- | --- | --- |
| *E. coli* |  |  |
| S17-1 | λpir | (7) |
| *B. bacteriovorus* |  |  |
| 109J | carrying pMQ581 | This study |
| *V. cholerae* |  |  |
| CNV 116 | N16961 *rbmA*-3xFLAG, *lacZ*:Ptac-mKate2 | (2) |
| CNV 121 | N16961 *rbmA*-3xFLAG, *lacZ*:Ptac-mKO-κ | (2) |
| CNV 127 | N16961, *lacZ*:Ptac-mKate2 Δ*rbmA* | This study |
| CNV 64 | *vpvC* W240R matrix hyper secretor, *lacZ*:Ptac-mKate2 | This study |
| CNV 252 | *vpvC* W240R matrix hyper secretor *rbmA*-3xFLAG, *lacZ*:Ptac-mKate2 | This study |

| Plasmid | Origin, marker | comments | source |
| --- | --- | --- | --- |
| pCN769 | pR6K, Amp | pBW *rbmA*-3xFLAG insertion | (2) |
| pMQ581 | RFS1010, Gent | Constructed by replacement of *tdTomato* with *gfpmut3* in pMQ414 parental plasmid (8) | This study |

## Supplemental Figures

**Figure S1:**
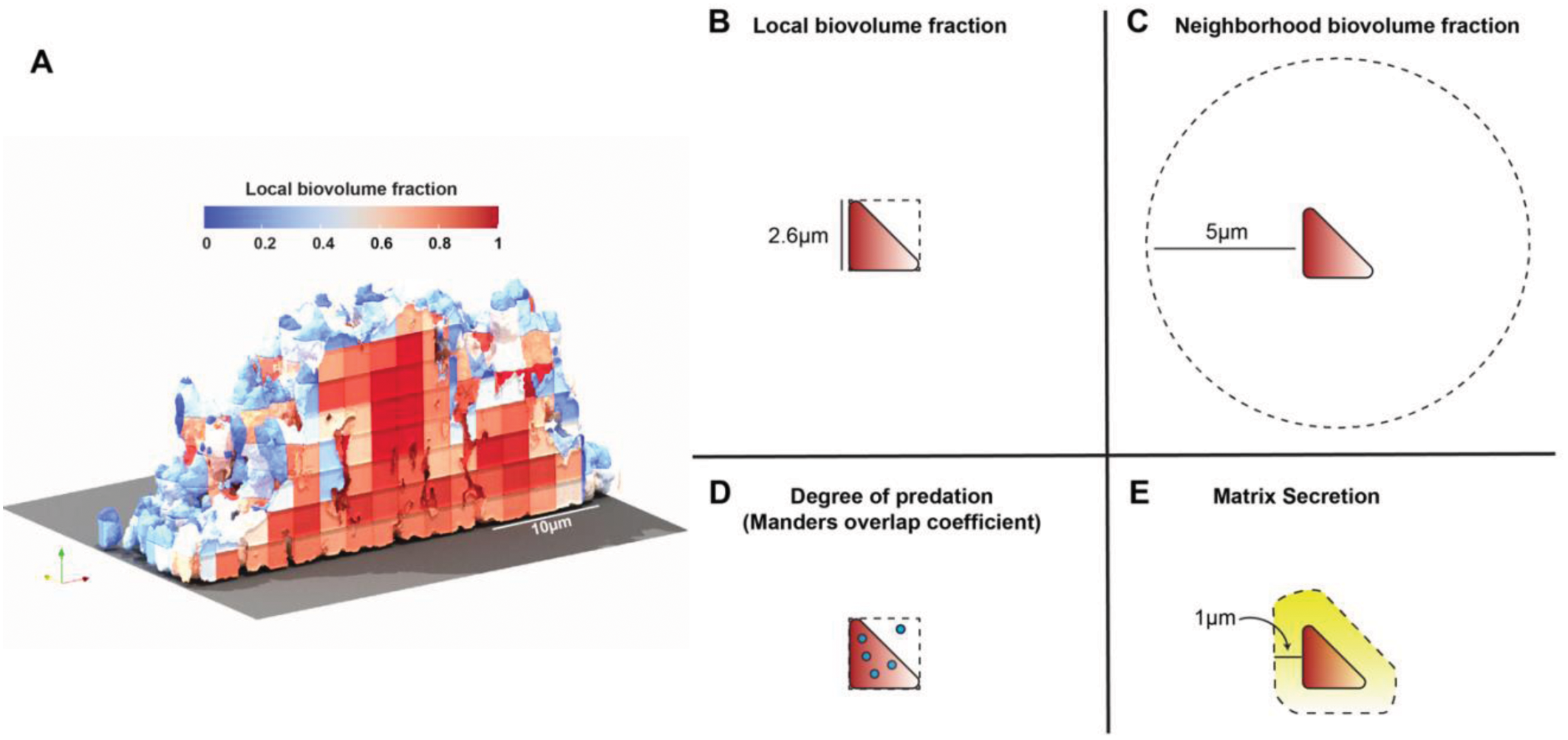
3D rendering of a segmented biovolume with cartoon representations of image analysis parameters. **(A)** 3D rendering how biofilms are segmented and dissected into a cubic grid for analysis. Grid unites here have been color-coded according to local biovolume fraction. (B) Local biovolume fraction measures the proportion of each grid unit occupied by *V. cholerae*. (C) Neighborhood biovolume fraction measures the proportion of the immediate neighborhood of each unit occupied by *V. cholerae*. (D) The extent of overlap (Manders overlap coefficient) describes the proportion of *B. bacteriovorus* signal that overlaps with *V. cholerae* signal within any given cube. We refer to this metric in the main text as ‘degree of predation’. (E) Matrix secretion measures signal intensity of matrix fluorescence within a 1µm shell around each segmented element of *V. cholerae*.

**Figure S2:**
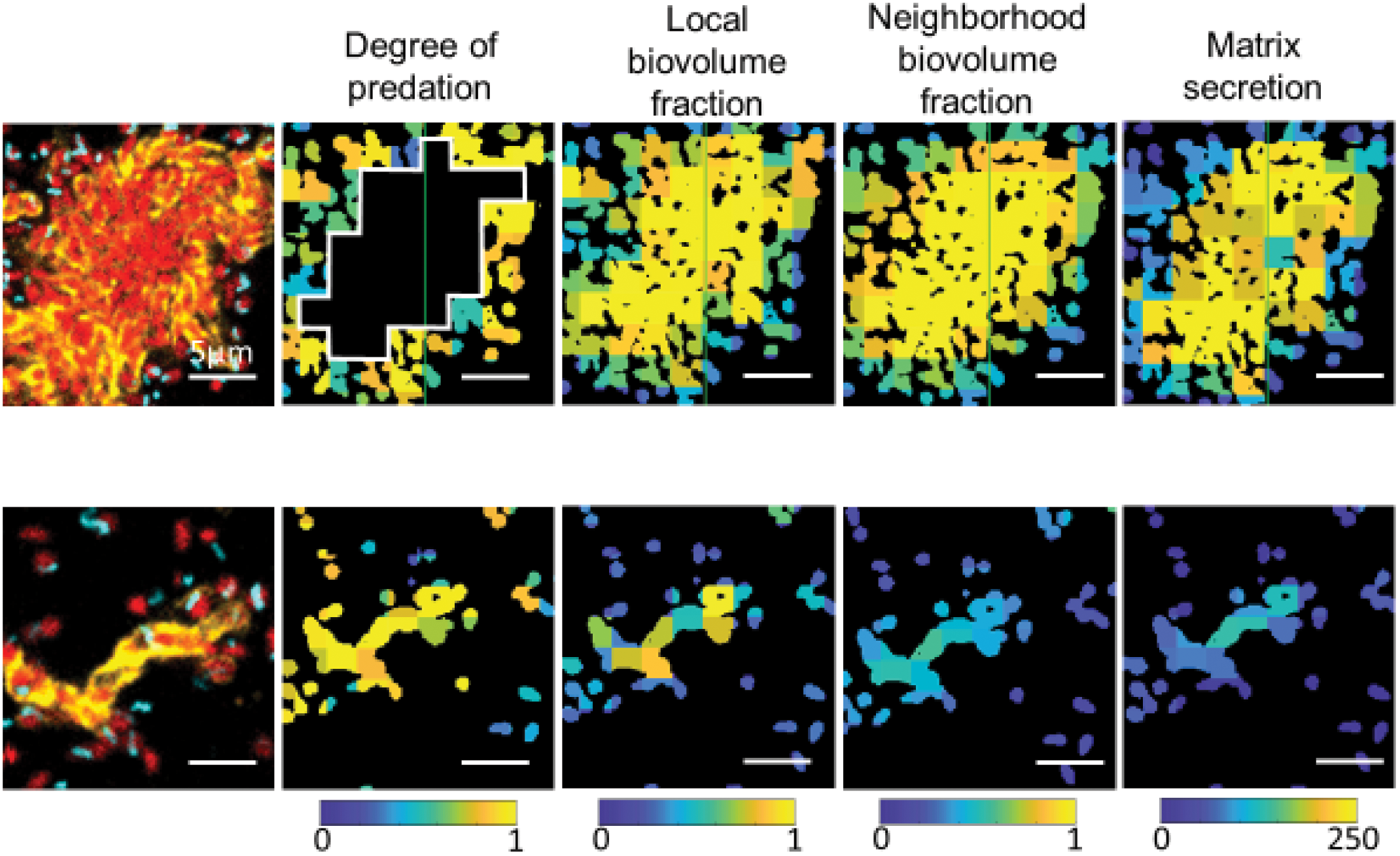
Heatmaps for each parameter of image cytometry analysis showing key differences between small and large matrix-positive cell clusters. Raw images are shown at left with degree of predation, local biovolume fraction, neighborhood volume fraction, and matrix secretion quantifications shown to the right.

**Figure S3:**
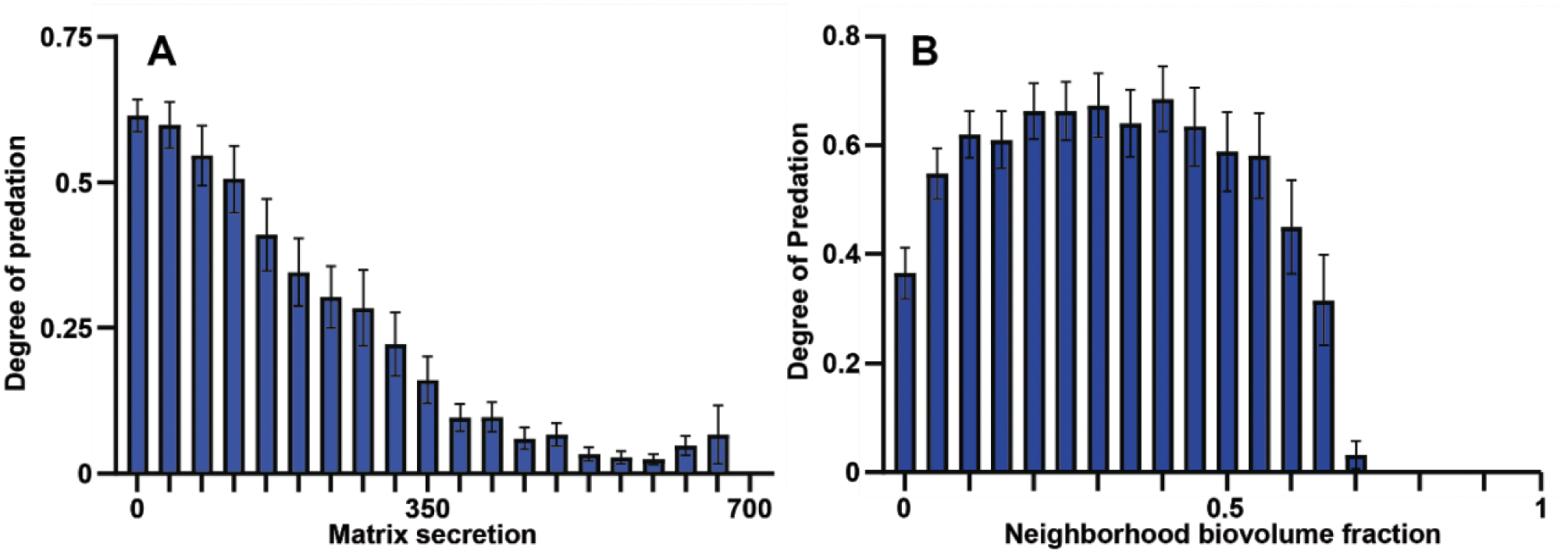
Contribution of matrix secretion and neighborhood biovolume faction to protection from *B. bacteriovorus* exposure. (A) Distribution of the degree of predation as a function of local matrix accumulation. (B) Distribution of the degree of predation as a function of neighborhood biovolume fraction (*n* = 23). The degree of predation decreases approximately linearly with local matrix accumulation, but decreases almost as a step-function as a function of neighborhood volume fraction.

**Figure S4:**
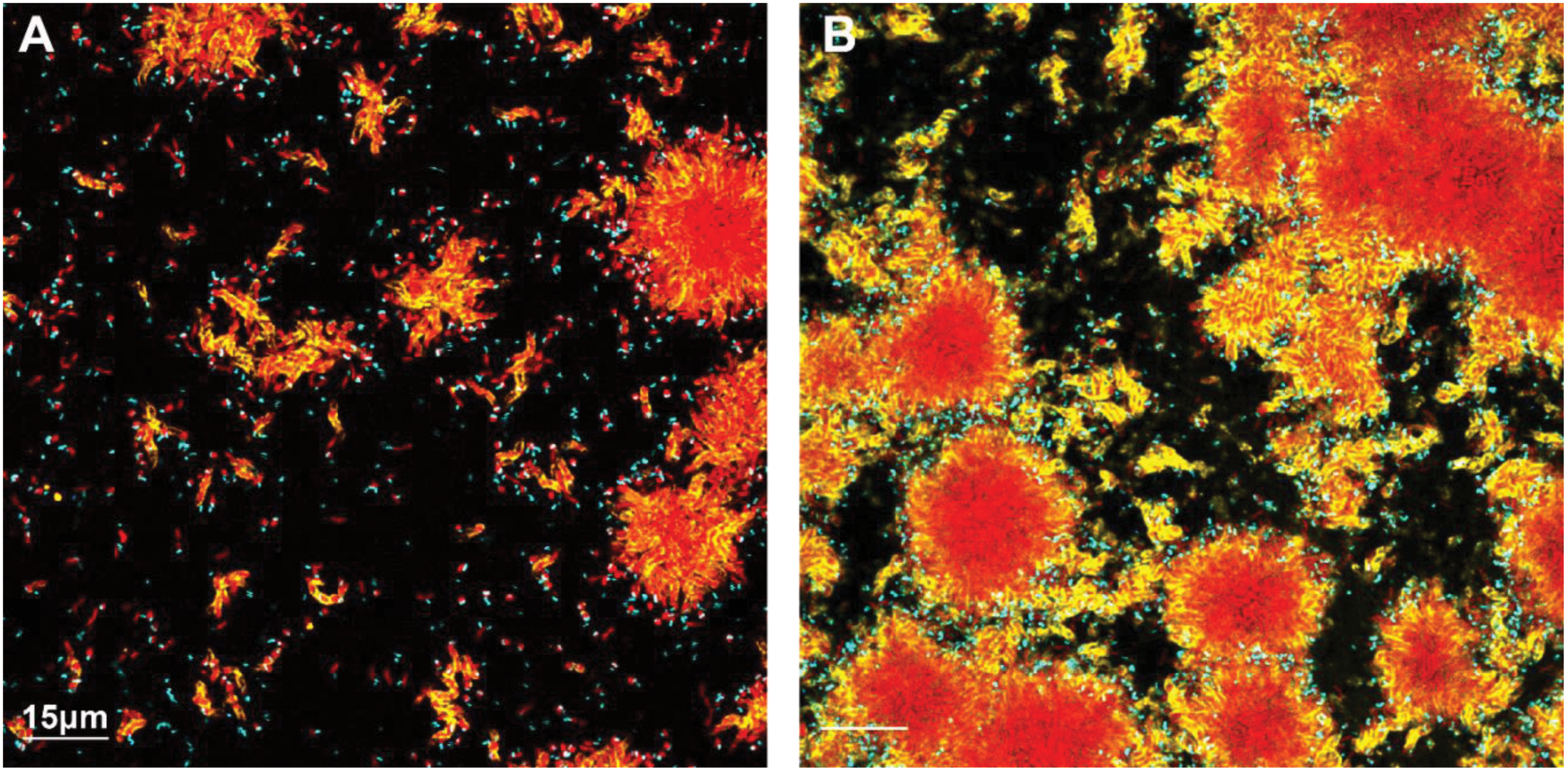
Wide view images of *V. cholerae* (A) wild type and (B) matrix hyper-secreting biofilms 2 hours after exposure to *B. bacteriovorus* predation. Resident *V. cholerae* biofilms are shown in red, biofilm matrix is shown in yellow, and *B. bacteriovorus* is shown in cyan.

**Figure S5:**
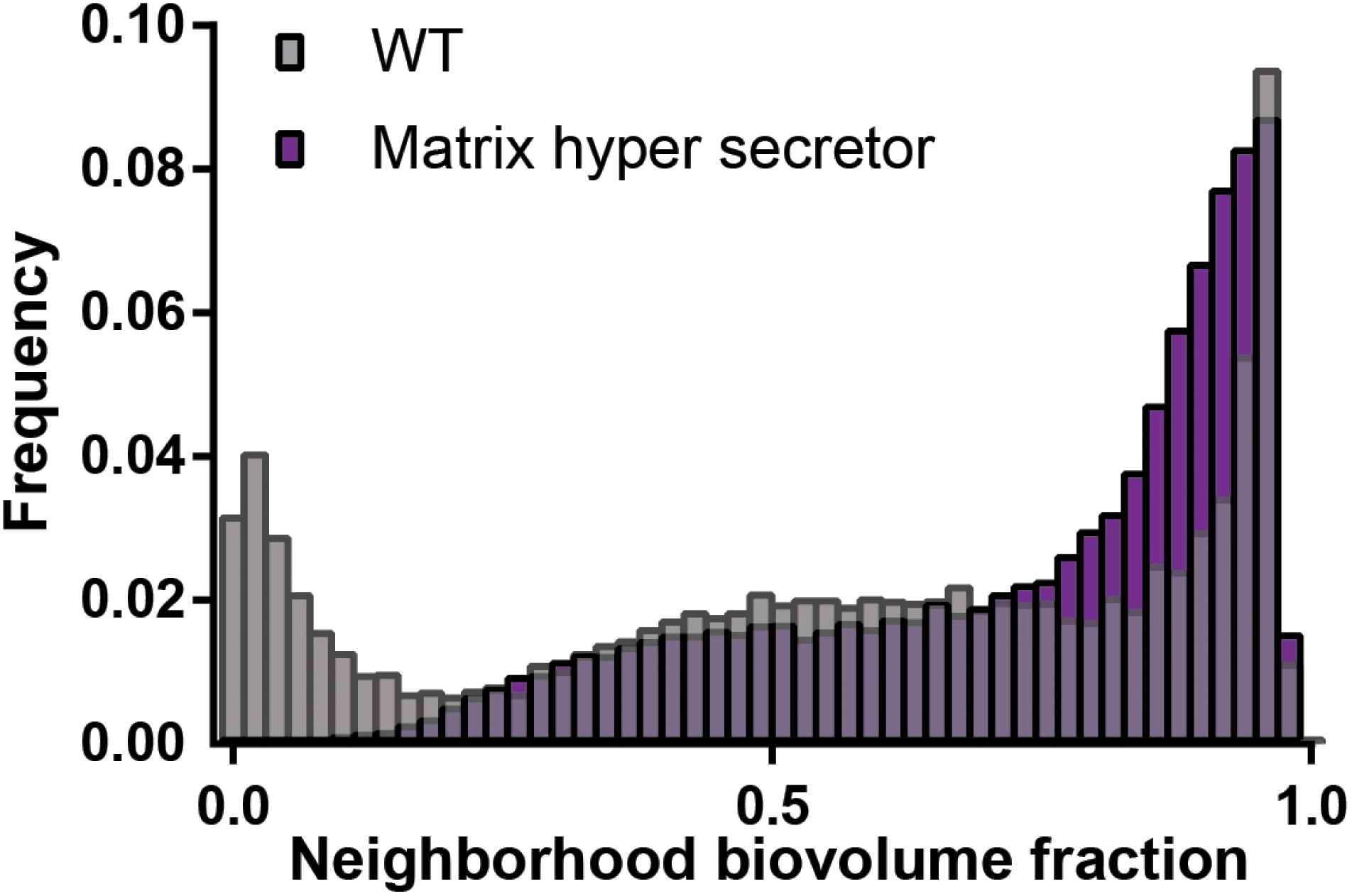
Neighborhood biovolume frequency distributions for biofilms of *V. cholerae* wild type (grey bars) and a matrix hyper-secretor variant (purple bars). Note that matrix hyper-secreting mutants have frequency distributions of cell-cell packing shifted toward higher values, allowing more biofilm clusters to survive. Both data sets were collected 2 hours post-exposure to predators (*n* = 9).

**Figure S6:**
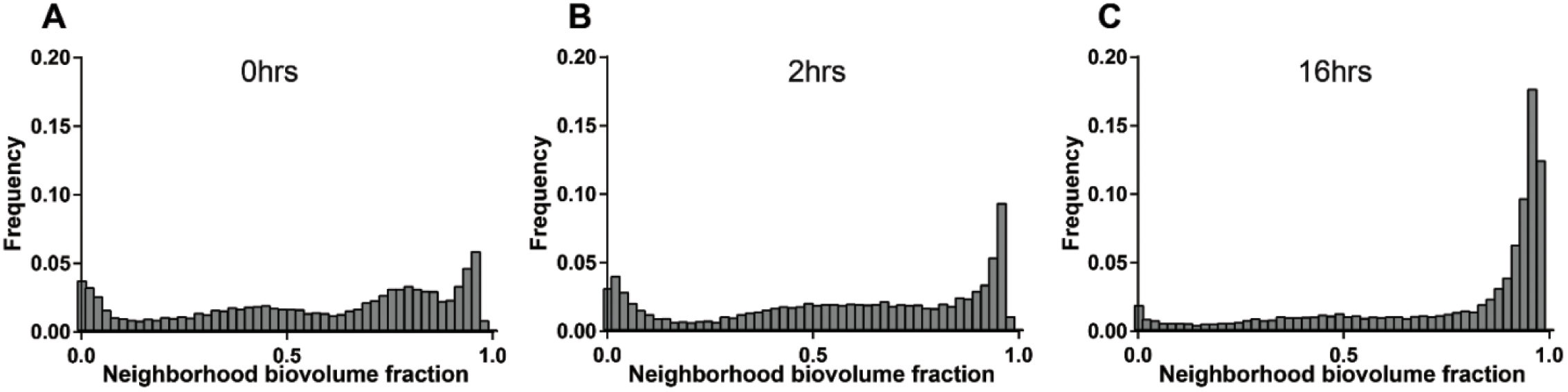
Time course reveals rapid change in cell packing density of resident *V. cholerae* wild type under *B. bacteriovorus* predation. Image stacks were taken at 3 time points following predator exposure (*n* = 15 per time point). While modest changes can be seen after 2 h of predator exposure, a large change can be seen after 16 hours, similar to the frequency diagram outlined in Figure 4C of the main text corresponding to biofilms 48 h after predator exposure.

**Figure S7:**
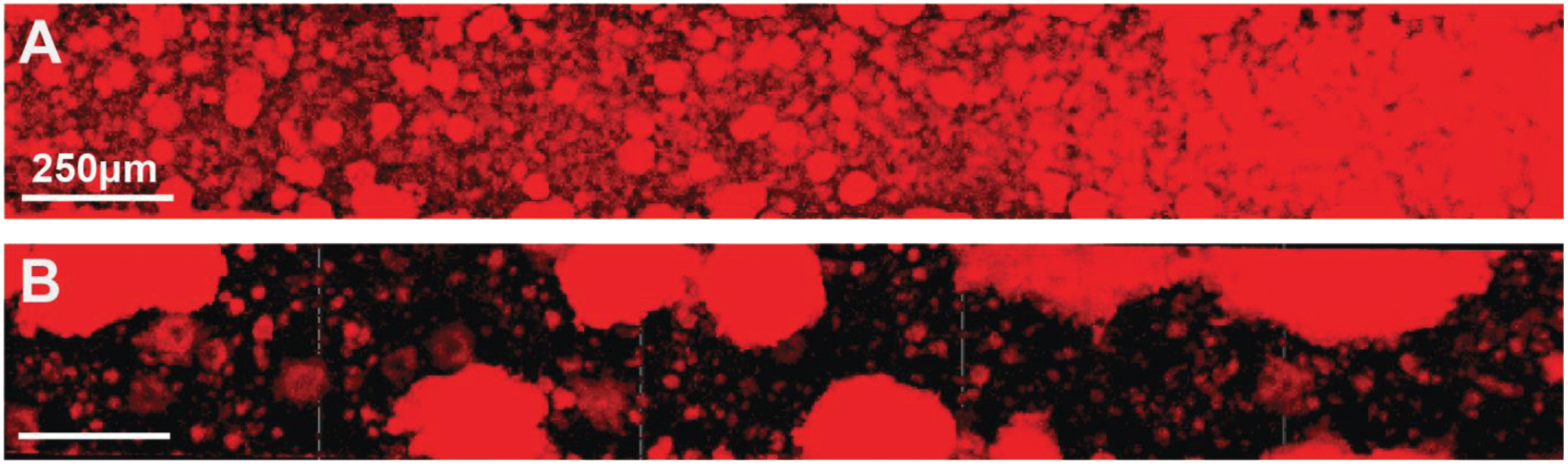
Submillimeter-scale landscape changes occur in *V. cholerae* biofilms following *B. bacteriovorus* predation pressure. View of an entire microfluidic device containing (A) *V. cholerae* un-exposed to predation or (B) *V. cholerae* biofilms 48 hours after predator exposure.

